# Interplay between the immune response and the adaptation of metabolic pathways upon infection

**DOI:** 10.64898/2026.01.02.697260

**Authors:** Andriy Goychuk, David Goh, Sergio Eraso, Ruslan Medzhitov, Arup K. Chakraborty

## Abstract

Glucose is the principal metabolic fuel for the energy needs of most cell types. Upon infection, cytokines secreted by the immune system regulate redistribution of glucose to meet new metabolic needs associated with clearing the pathogen. We develop a mathematical model to describe the dynamics of such adaptation of metabolic pathways mediated by the immune response and its impact on the ability to clear pathogen and restore health. We find that cytokine-regulated redistribution of glucose resources in different tissues is critical for an effective immune response to pathogen as strictly clamping plasma glucose levels to homeostatic levels results in an ineffective immune response. By studying the effects of various parameters in our model, we describe how aberrant regulation of adaptation mechanisms affect outcomes of infection. Too high a glucose consumption rate by innate immune cells to mediate functions results in failure to clear pathogen. Pathogens with a very high replication rate can be controlled to low levels, but at a very high metabolic cost. Too low a pathogen replication rate allows the pathogen to hide from the immune system and rebound to high levels at later times. Finally, the strength of the innate immune response must be regulated to not be too high, not only to limit immunopathogenesis, but also for mediating an effective adaptive immune response.

## Introduction

The energy needs of different physiological systems are met by metabolic fluxes that are tightly regulated by hormones. One key metabolite is glucose, which is the main metabolic fuel for ATP production in most cell types, as well as a source of carbon for anabolic processes. The systemic glucose concentration is regulated by insulin and glucagon and maintained around the homeostatic set point of 5 mM in healthy adults. Cells take up glucose through several GLUT family glucose transporters that are characterized by tissue-specific expression, different modes of regulation, and Michaelis constants *K*_*m*_. GLUT1 (often reported *K*_*m*_ ~ 3 mM to 7 mM [1, 2], with some studies reporting *K*_*m*_ ~ 1 mM to 2 mM [3, 4]) is used by most cell types including immune cells; GLUT2 (reported *K*_*m*_ ~ 17 mM to 20 mM [1, 2, 3]) is a bidirectional transporter expressed in liver, intestine, kidney, and pancreatic beta cells; GLUT3 (reported *K*_*m*_ ~ 1 mM to 2 mM [2, 3, 4]) is expressed in neurons; and GLUT4 (reported *K*_*m*_ ~ 5 mM [2, 3] but also as high as *K*_*m*_ ~ 6.6 mM to 8.2 mM [1, 4]) is expressed in muscle and adipose tissue. The high glucose affinity of GLUT3 enables neurons to take up glucose even under hypoglycemic conditions and thus insulates the brain from daily fluctuations in glucose levels. The low affinity of GLUT2 is utilized for sensing blood glucose concentration by beta cells in rodents (whereas human beta cells mostly use the higher-affinity GLUT1), for storing glucose surplus by hepatocytes, and for secretion of endogenously synthesized glucose by the liver and kidneys. In contrast, GLUT1 and GLUT4 establish regulated glucose uptake, such that plasma membrane expression of these transporters is controlled either by external signals (insulin, growth factors, and cytokines) or by the cell-intrinsic metabolic demands. Specifically, under conditions of glucose surplus, the insulin signal, through the kinase AKT, induces GLUT4 expression in skeletal muscle and adipose tissue, leading to glucose utilization or storage in the form of glycogen or lipids, respectively. In addition, when skeletal muscle glucose is depleted during exercise, GLUT4 expression is promoted by the ATP sensor kinase AMPK. Similarly, GLUT1 expression in most cell types is induced by AKT in response to growth factors or cytokines, or by AMPK in response to high metabolic demand. In addition, inflammatory cytokines suppress insulin-induced GLUT4 expression and GLUT2-dependent insulin secretion, thus preventing glucose storage and making it available for the immune system, where cytokines promote GLUT1 expression.

Given the essential role of glucose homeostasis in most physiological processes and the critical role of glucose as a metabolic fuel for the immune system, here we seek to develop an understanding of the dynamics of the processes by which glucose regulation is modulated by immune responses, and thus also obtain insights into how this interplay can be aberrantly regulated. Previous computational work by Zhao et al. [5] focused on the crosstalk between the brain, the immune system, energy storage in fat, and energy supply through feeding, by employing a coarse-grained energy flux and food uptake model. Here, we explicitly account for the effects of cytokine signaling in the regulation of how glucose is redistributed in physiological systems upon infection. We calibrate glucose fluxes at homeostasis to experimental data. Our main findings include that cytokine-mediated regulation of glucose uptake in tissues and immune cells is essential for an effective host immune response. However, counterintuitively, an overactive innate immune response can be detrimental to the host, not just because of immunopathogenesis, but also because the resulting low pathogen load leads to a weak adaptive immune response. Moreover, our model predicts that plasma glucose levels can either increase or decrease depending on the severity of the infection (represented by pathogen replication rate). Taken together, our results provide guides for future experiments that probe the interplay between the immune response and adaptation of metabolic pathways upon infection. Some of our results may also apply to situations where the interplay between the adaptation of a limiting resource and the immune response upon physiological perturbations is important for maintaining a healthy or diseased state.

## A model for regulation of glucose fluxes to physiological systems upon infection

Figure 1 depicts the regulatory network that maintains glucose homeostasis and how it is modulated by cytokines upon infection, as well as how the immune system acts on replicating pathogens. Below, we describe our model for different components of this regulatory network.

**Fig. 1:**
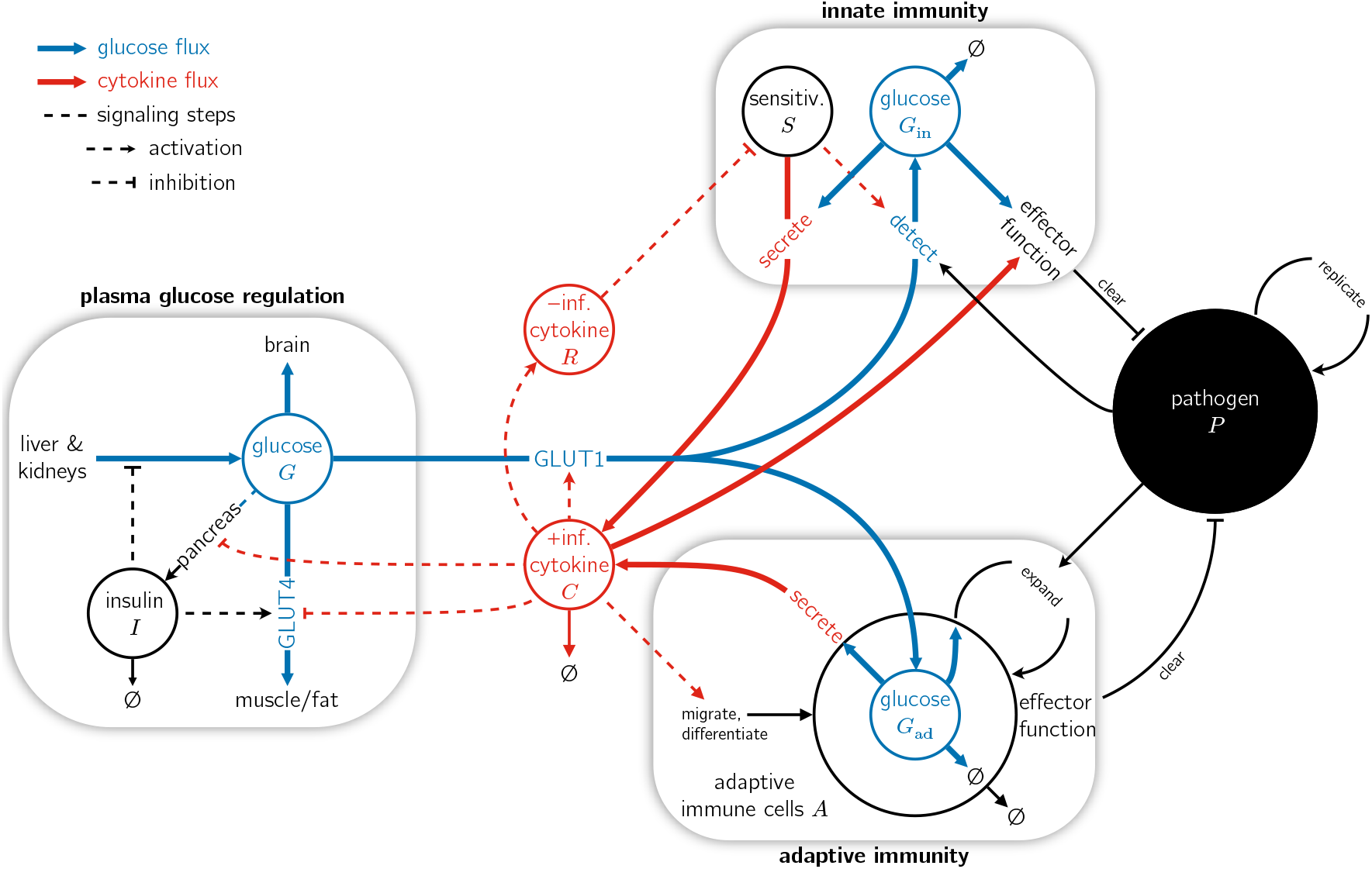
Schematic depicting the regulatory network that mediates glucose fluxes to tissues and how the immune response upon infection modulates it. Cytokines secreted by immune cells redistribute resources in the form of glucose among different tissues and immune cells. Glucose fluxes are depicted by blue arrows, whereas cytokine fluxes are shown by red arrows. Dashed lines depict regulation by cytokines, with bar caps indicating inhibition and arrow caps indicating activation. Black arrows correspond to processes that affect pathogens or vice versa. +inf. refers to inflammatory cytokines and −inf. to anti-inflammatory cytokines. *S* is the sensitivity of the innate immune cells in detecting pathogen.

### Glucose and insulin regulation

In the postprandial state (after a meal), glucose is secreted primarily by the intestine, and the liver stores glucose in the form of glycogen [6, 7]. To minimize the complexity of our mathematical model and the number of unknown parameters, we developed a description tailored to the nighttime fasting state when the immune response is strongest [8, 9]. We neglect other effects of the circadian rhythm such as ingestion of meals, etc., during infection. In doing so, we lump all glucose producers into a single source, namely the liver which secretes glucose.

### System-wide glucose flux balance

We begin by accounting for all glucose fluxes that enter and leave the bloodstream. We explicitly model the liver, brain, and muscle/fat as the main tissues that produce and consume glucose in homeostasis. We also explicitly consider the immune response upon infection, and its effects on the pathogen and metabolic fluxes to different organs. The generation of immune responses consumes energy in the form of glucose. We model the glucose dynamics upon infection as follows:

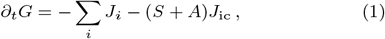

The quantity on the left in Eq. (1) is the time derivative of the plasma glucose concentration (*G*). The glucose uptake by the innate immune system is proportional to *S*, which is a measure of its sensitivity in detecting pathogen. The glucose uptake by the adaptive immune system is proportional to the number of recruited cells of the adaptive immune system, whose total biomass we represent with the variable *A*. The sensitivity, *S*, decreases with the duration of inflammation, whereas the biomass of the adaptive immune system *A* increases and then declines as the host battles and clears infection. The variable *J*_ic_ models the cytokine-regulated glucose flux into immune cells. The total flux increases with the biomass of the adaptive immune system or with the sensitivity of the innate immune system. Finally, *J*_*i*_ is the flux of glucose into each tissue, *i* ∈ {liver, brain, muscle/fat}. We normalize the plasma glucose concentration *G* by its homeostatic value *G*_s_ = 5 mM as it is a natural measure [10].

In the following, we detail the different terms in Eq. (1) that describe glucose fluxes into each tissue and how they are regulated by the immune response, as well as glucose consumption by the immune system. These glucose fluxes are represented by thick blue arrows in Figure 1.

### Hepatic glucose production

During fasting, glucose is secreted into plasma by two sources, namely the liver and kidneys, which account for ~ 75% and ~ 25% of total glucose production, respectively [11]. The substrates for glucose production are glycogen stored in the liver (glycogenolysis) and metabolites circulating in the blood (gluconeogenesis) [6]. Insulin suppresses glucose secretion by the kidneys and the liver with approximately equal efficacy [12, 11]. In Fig. 1, this is depicted by a dashed line with a bar cap. In our model, we rely on measurements of the hepatic glucose production rate as a function of insulin concentration in humans [13]. These experimental data show that hepatic glucose production decreases with increasing concentration of the hormone insulin [Fig. 2A] and can be fitted with [14]

**Fig. 2:**
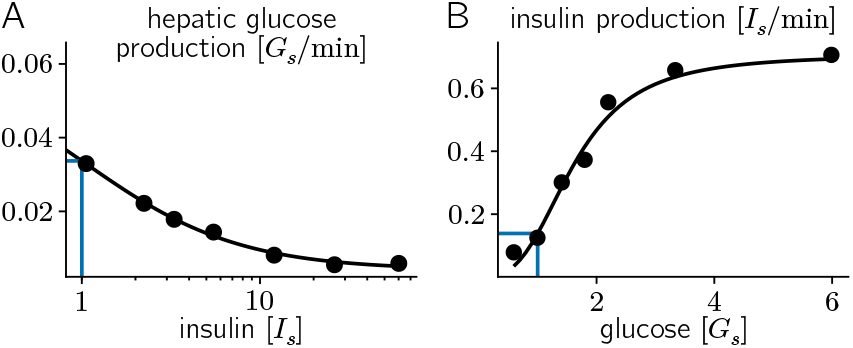
A) Glucose production by the human liver as a function of plasma insulin concentration. Data digitized from Ref. [13] and reused with permission from Elsevier (license 6172150320873). Solid black line corresponds to best fit of Eq. (2). Blue lines indicate typical values in homeostasis. B) Insulin production by beta cells in the pancreas as a function of plasma glucose concentration. Data digitized from Ref. [16] and reused under Creative Commons Attribution License (CC BY). Solid black line corresponds to approximation of best fit reported in Ref. [16], as discussed in the main text. Blue lines indicate typical values in homeostasis.

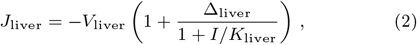

where the sign indicates an export from the liver into the blood plasma.

We normalize the plasma insulin concentration *I* by the typical value *I*_s_ = 30 pM as it is a natural measure [10]. The best-fit parameters in Eq. (2) are *V*_liver_ = 4.1 × 10^−3^ *G*_s_*/*min for the maximal rate of glucose efflux from the liver, *K*_liver_ = 1.0 *I*_s_ for the inhibitory constant, and Δ_liver_ = 14.1 for the relative amount of insulin-inhibited hepatic glucose production [Fig. 2A]. The hepatic glucose production rate predicted by Eq. (2) in homeostasis, 3.3×10^−2^ *G*_s_*/*min, compares favorably with reported values of the endogenous glucose production rate, 3.2 × 10^−2^ *G*_s_*/*min, by Basu et al. [10]. Glucose secretion is also stimulated by the hormone glucagon [15], which is antagonistic with insulin and coarse-grained in the effective fitting parameters of Eq. (2). The effect of pro-inflammatory cytokines on insulin, and hence *J*_liver_ is described below.

### Pancreatic insulin secretion

In non-diabetic individuals, insulin is secreted by beta cells in the pancreas in response to elevated blood glucose levels. The rate of insulin secretion has been measured experimentally [16] and follows a Hill curve as a function of plasma glucose [Fig. 2B]. Together with insulin degradation by different tissues, this leads to the following plasma insulin dynamics:

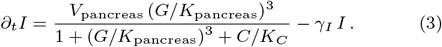

The Hill coefficient of approximately three is based on *in vitro* data reported in Ref. [16], and is higher than the value of two cited by Ref. [17]. However, the experimental data shown in Fig. 2B can be well approximated by either choice. The effective Michaelis constant, normalized to the homeostatic glucose concentration, is given by the best-fit *K*_pancreas_ = 1.6 *G*_s_ reported in Ref. [16]. This is higher than the Michaelis constant of GLUT1 [1, 2, 3, 4], which facilitates glucose transport in human beta cells [18]. Hence, glucose uptake is potentially not the only rate-limiting step for insulin production. The rate of insulin degradation according to Ref. [17] is *γ*_*I*_ = 0.14*/*min, and the maximum rate of insulin secretion *V*_pancreas_ = 0.71 *I*_s_*/*min is chosen to match typical plasma insulin concentrations in homeostasis.

In Eq. (3) we have taken into account that pro-inflammatory cytokines inhibit glucose-stimulated insulin secretion [19, 20]. The category of pro-inflammatory cytokines refers to many different biomolecules that are present at different concentrations. Therefore, we normalize the concentration of pro-inflammatory cytokines by an unspecified effective value, *C*_*s*_, which serves as a natural measure. It is not possible to determine the inhibitory constant *K*_*C*_ from experimental data, because cytokine concentrations are extremely small, and it is therefore a free parameter. For our simulations, we choose its value as follows. For small values of *K*_*C*_ ≪ *C*_*s*_, insulin secretion is always suppressed regardless of the cytokine level *C*, which is not consistent with physiology. For large values of *K*_*C*_ ≫ *C*_*s*_, cytokines have no effect at physiological concentrations, which is also inconsistent with experimental evidence [19, 20]. Therefore, we choose *K*_*C*_ ≡ 1 ×*C*_*s*_ to make insulin secretion sensitive to cytokines.

Because cytokines inhibit insulin secretion [Eq. (3)], and insulin inhibits hepatic glucose production [Eq. (2)], it follows that cytokines can indirectly stimulate hepatic glucose production. Pro-inflammatory cytokines have also been reported to directly stimulate gluconeogenesis in the liver [21], which we have neglected here.

### Insulin-dependent glucose uptake

In addition to regulating hepatic glucose production, insulin also controls the amount of glucose transporters on the membranes of muscle and fat cells, and thereby regulates their glucose uptake [22, 17]:

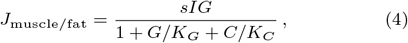

where the sign indicates an influx into muscle and fat tissue. Note that there is no backflow of glucose, assuming that glucose is quickly phosphorylated by hexokinase once it enters the cell [23]. The glucose transport [Eq. (4)] follows Michaelis-Menten kinetics with the apparent dissociation constant *K*_*G*_ ≈ 2 *G*_s_ determined by experimental data on human muscle cells and the insulin-dependent transporter GLUT4 [24, 4]. The insulin sensitivity, *s* = 2.6 × 10^−2^*/*(*I*_s_ min), is numerically optimized such that at homeostatic insulin concentrations *I* = *I*_*s*_, and for the cytokine concentration in the absence of pathogen, steady state plasma glucose levels are controlled at *G* = *G*_s_.

Note that in Eq. (4), an increase in the cytokine concentration *C* will decrease the effective insulin sensitivity, which is the inverse of glucose resistance. This takes into account that pro-inflammatory cytokines such as TNF-*α* induce insulin resistance by interfering with insulin signaling and reducing the expression of insulin-dependent glucose transporters [25, 26, 27, 28]. Thus, Eq. (4) models the regulation of glucose flux into muscle/fat by cytokines (depicted by a dashed line with a bar cap in Fig. 1). Using similar arguments as employed above, and the lack of experimental measurements, we assume that the (effective) inhibitory constant in Eq. (4) is *K*_*C*_ ≡ 1 × *C*_*s*_ to make glucose uptake sensitive to cytokines. In future studies, one could use different values for the inhibitory constants *K*_*C*_ in Eqs. (4) and (3) to study scenarios in which either insulin secretion or insulin-dependent glucose uptake are more sensitive to cytokines. Note that, together with the inhibition of glucose-stimulated insulin secretion by pro-inflammatory cytokines [19, 20], which we discussed in the preceding section, cytokine-induced inhibition of glucose uptake by muscle/fat will increase plasma glucose levels.

Interestingly, there are also reports that TNF-*α*, secreted by muscle cells under stress, can increase the amount of glucose transporters on the cell surface via autocrine and paracrine signaling [29]. Hence, glucose uptake into muscle and fat cells could be a nonmonotonic function of cytokine concentration, which points to the multifaceted action of cytokines in homeostatic control [30]. We will neglect such effects because we assume that muscle is not always under stress upon infection.

### Insulin-independent glucose uptake

Among the various different glucose transporters, GLUT4 is the main transporter whose abundance in the plasma membrane is regulated by insulin. Conversely, this means that there are many tissues in the body, such as the liver, pancreas, heart and brain, that consume glucose independently of insulin. Out of these organs, the liver acts as a glucose source during fasting and the pancreas is a negligible consumer—its main contribution to glucose regulation is taken into account by insulin secretion according to Eq. (3). Therefore, we will now focus on the brain, which is the organ that consumes the most glucose.

The concentration *G*_brain_ of glucose in the brain of human volunteers, in response to clamping their plasma glucose levels *G*, have been measured with nuclear magnetic resonance in Ref. [31]. Here, we fit these experimental data with a simple linear model, *∂*_*t*_*G*_brain_ = *J*_brain_ − *γ*_brain_*G*_brain_, where the net glucose flux is given by [Fig. 3]

**Fig. 3:**
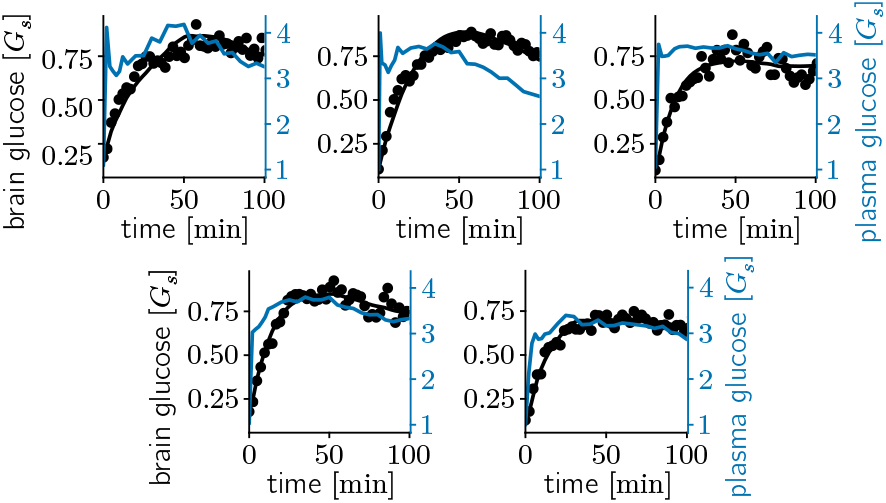
Dynamics of glucose concentration in the brain as a function of plasma glucose levels. Experimental data was obtained from the authors of Ref. [31]. Solid black lines correspond to best-fit of the phenomenological brain dynamics [Eq. (5)] to the experimental data (black dots). To fit the brain glucose dynamics as only degree of freedom, we interpolated the empirical measurements of the plasma glucose concentration as a function of time (solid blue lines) and used this as input in the fitting procedure.

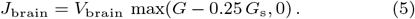

This phenomenological model is chosen to reproduce the linear relationship between blood and brain glucose levels seen in experiment [32, 31]. This linear relationship has its origins in the reversible transport of glucose across the blood brain barrier [32]. In Eq. (5), we have assumed that the brain only takes up glucose if the plasma concentration exceeds roughly 0.25 *G*_s_ = 1.25 mM. This is motivated by the extrapolation of experimental data indicating that there is only glucose in the brain if the plasma glucose concentration exceeds a threshold value [32, 31]. This threshold value does not need to be very high because glucose transport is dominated by GLUT1 for crossing the blood brain barrier and GLUT3 for uptake into neurons, two high-affinity transporters that have similar Michaelis constants as the above threshold [33].

Because brain glucose levels simply track plasma glucose levels according to Eq. (5), we do not have to monitor the dynamics of brain glucose levels separately. The best-fit parameters in Eq. (5) are *V*_brain_ = 2.1 × 10^−2^*/*min for the uptake rate and *γ*_brain_ = 9.0 × 10^−2^*/*min for glucose turnover in the brain. These values correspond to a glucose uptake of roughly 420 kcal*/*d or 20% of the total metabolic needs of the body, in agreement with the existing literature [34, 35].

### Immune reactions and metabolism

#### Pathogen replication and neutralization

There are three classes of replicating pathogens that can infect a host: (a) viruses, (b) microorganisms such as bacteria, amoeba, or fungi, and (c) multicellular parasites. Each class has a different mode of replication and elicits a different immune response. For example, viral replication has been described in the past literature with different variants of the target-infected-virus (TIV) model [36] which is conceptually similar to the susceptible-infected-recovered (SIR) model used in epidemiology [37]. Although the details of infections with different types of pathogen may be different, the phenomenological dynamics are similar: pathogen *P* replicates with rate *k*_infect_,

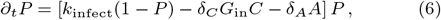

and is cleared by the innate immune response, with rate *δ*_*C*_ *G*_in_*C*, and by the adaptive immune response, with rate *δ*_*A*_*A*. The clearance of pathogen by the innate immune response, with rate constant *δ*_*C*_, depends on the availability of glucose *G*_in_ to innate immune cells due to the reliance on glycolysis for their energy supply [38]. We take the clearance rate to be proportional to the concentration (*C*) of pro-inflammatory cytokines as a proxy for the many effects of innate immunity such as macrophage, neutrophil, eosinophil, and basophil activity. The clearance of pathogen by the adaptive immune response depends on its biomass *A*, which subsumes effector T cells and antibodies. Thus, while the expansion of adaptive immunity does require glucose as will be discussed next, with glycolysis providing the metabolites required for generating biomass, we assume that pathogen clearance by adaptive immune cells (e.g., by degranulation of cytotoxic T cells) is metabolically inexpensive in comparison and hence not rate-limited by glucose availability.

Note that we have normalized the concentration of pathogen by its carrying capacity, so that *P* = 1 corresponds to all target cells being infected. For pathogen to be eliminated, *δ*_*C*_*G*_in_*C* +*δ*_*A*_*A* must exceed the pathogen replication rate. We initially set *δ*_*C*_ = 0.001*/*(*C*_s_ *G*_s_ min) for the rate of cytokine-dependent pathogen clearance and *δ*_*A*_ = 0.005*/*min for the rate of pathogen clearance by adaptive immunity, but will vary these rate constants later and examine the effects. For the rate of pathogen replication, we initially set *k*_infect_ = 1.0 × 10^−3^*/*min and will also study variations in its value. These rates are chosen such that infection and clearance proceed on a typical timescale of about one day. To fully recover from infection, the innate and adaptive immune responses should remain engaged long enough to push pathogen levels *P* below the extinction threshold. We choose the extinction threshold as *P*_clear_ = 10^−4^ which is one order of magnitude smaller than the levels of pathogen at the time of infection, the latter being chosen as *P* (*t* = 0) = 10^−3^.

### Cytokine-dependent activation and expansion of adaptive immunity

To model the initiation and activation of adaptive immunity by the innate immune response, we consider a cytokine-dependent influx and differentiation of adaptive immune cells into effector phenotypes [39], with rate

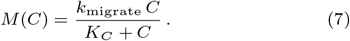

As before [cf. Eqs. (3) and (4)], because we want the rate of differentiation to be sensitive to the cytokine levels, we set *K*_*C*_ ≡ 1 × *C*_*s*_. Note that one should not choose the parameter *k*_migrate_ to be too high, otherwise a positive feedback loop is set up because adaptive immune cells secreting cytokines (discussed below) will recruit even more adaptive immune cells, which can keep the adaptive immune response active even when pathogen has been cleared. Thus, we set the rate constant to *k*_migrate_ = 9.6 × 10^−5^*/*min so that the adaptive immune response would take multiple days to fully expand, unless we take into account the replication of adaptive immune cells, as will be discussed next.

Previous work has studied the expansion of T cells when they bind to pathogen-derived antigen [40], with the duplication rate given by

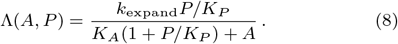

We use this as a proxy for all adaptive immune cells and choose the free parameter *K*_*P*_ = 0.01 to be small so that adaptive immune cells can recognize pathogen at low concentrations. The adaptive immune system needs to expand substantially to eliminate pathogen. Therefore, to allow large carrying capacities, we choose a large inhibitory constant *K*_*A*_ = 0.1. We set the rate constant of adaptive immune system expansion to *k*_expand_ = 1.6 × 10^−4^*/*(min *G*_s_), which corresponds to a timescale of multiple days at homeostatic glucose levels.

Finally, building on these ideas, we also assume that glucose *G*_ad_ in adaptive immune cells is a key resource required by activated, proliferating immune cells [41, 42, 43, 44]. Taken together, we model the dynamics of the adaptive immune system as follows:

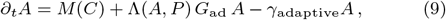

where *γ*_adaptive_ = 2.4 × 10^−4^*/*min is the rate of inactivation of adaptive immunity. We have chosen this rate constant to be large enough so that the immune response ceases within a few days after pathogen has been cleared. At the same time, this rate constant cannot be too large, lest the immune response ceases before pathogen has been fully eliminated.

### Cytokine secretion by innate and adaptive immunity

Immune cells secrete cytokines in response to infection. Cytokines can rewire the glucose regulatory circuit as described in Figure 1 and Secs. 2.1.3 and 2.1.4. We assume that cytokine secretion by innate immunity depends on the presence of pathogen *P* and the sensitivity *S* of innate immune cells to pathogen. At the beginning of our simulations, we set *S*(*t* = 0) = 1 so that innate immune cells are primed to detect invading pathogen, and then reduce the sensitivity over time to prevent excessive immunopathology, as will be described next.

Inflammation due to pro-inflammatory cytokines *C* and the resulting activity of effector cells can lead to excessive tissue damage and is therefore regulated by several antagonistic mechanisms such as the secretion of anti-inflammatory cytokines [45] or changes in transcription due to tissue acidification [46]. Here, we abstract these regulatory mechanisms into a single variable *R*, which increases with prolonged inflammation and controls the sensitivity of innate immune cells with respect to pathogen via negative feedback:

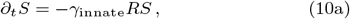

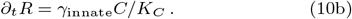

We choose the desensitization rate, *γ*_innate_ = 3.2×10^−4^*/*min, such that the innate immune response begins to wane, and the adaptive immune response takes over on a timescale of a few days.

Adaptive immune cells like CD4 T cells also secrete cytokines. We thus model the dynamics of pro-inflammatory cytokines, with concentration *C*, as follows:

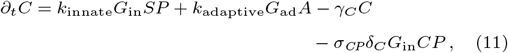

where we assume that cytokine secretion is conditioned on the glucose supply of innate and adaptive immune cells [42, 47, 48]. For the degradation rate of cytokines, *γ*_*C*_, since previous literature [49, 50] has reported half-lives of 20 min to 30 min in plasma depending on the cytokine, we choose *γ*_*C*_ = 3.5 × 10^−2^*/*min. The last term in Eq. (11) models the rate of cytokine consumption by effector cells that neutralize pathogen and is proportional to the corresponding killing rate [Eq. (6)]. We will first neglect this term by setting the amount of cytokines consumed per unit of pathogen cleared to *σ*_*CP*_ = 0, and will later explicitly study its effect on the dynamics.

The remaining parameters are chosen on the following basis. For the rate of cytokine secretion by innate immunity, we choose *k*_innate_ = 0.17*/*(*G*_s_ min) so that the corresponding timescale is short (taken to be similar to the timescale corresponding to insulin degradation); this parameter only sets the level of cytokines that will ultimately be reached. For the rate of cytokine secretion by adaptive immunity, we choose *k*_adaptive_ = 0.09*/*(*G*_s_ min). This parameter choice is constrained by the following considerations: Very large values of *k*_adaptive_ lead to a feedback loop where the cytokine-dependent activation of adaptive immune cells [cf. Eq. (7)], coupled with their cytokine production, leads to bistability and an immune response that persists even after pathogen has been cleared. Very small values of *k*_adaptive_ are insufficient for an appreciable expansion of adaptive immunity [see Eqs. (7) and (9)] and thus make the immune response to pathogen inadequate. Thus, an effective immune response requires intermediate values of *k*_adaptive_.

### Cytokine-dependent glucose uptake and homeostasis

In this section, we connect the secretion of cytokines with metabolic control. Pro-inflammatory cytokines stimulate glucose uptake by immune cells through the glucose transporter GLUT1 [51, 47, 52]. We model the corresponding glucose flux as

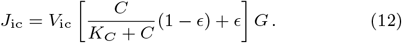

Here, the idea is that glucose uptake is small as long as the concentration of pro-inflammatory cytokines is small, *J*_ic_ ≈ *ϵV*_ic_*G* with *ϵ <* 1. When cytokines are released in response to infection, glucose uptake by immune cells increases due to GLUT1 expression [51], up to a maximum glucose flux of *J*_ic_ ≈ *V*_ic_*G*. As before, the sign indicates that glucose flows from the plasma into the immune compartment. The parameters are not known experimentally, and we choose *ϵ* = 0.25 and *V*_ic_ = 0.02*/*min. The rationale underlying this choice is that immune cells could metabolically starve if *ϵ* is too small, whereas for *ϵ* ~ 1 there would be no regulation of glucose flux by cytokines. The maximum uptake rate *V*_ic_ is chosen to roughly match the glucose consumption of other organs.

Next, we will discuss the glucose dynamics in the innate and adaptive immune compartments. We assume that each immune cell has a baseline glucose consumption rate to maintain its basic functions. We choose this baseline glucose consumption rate so that the steady-state glucose concentration is similar to the plasma glucose concentration for large cytokine concentrations and is very small for small cytokine concentrations. For the innate immune compartment, we model the glucose dynamics as follows:

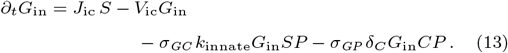

Here, *σ*_*GC*_ controls how much glucose innate immune cells use to produce cytokines and *σ*_*GP*_ how much glucose effector cells use to neutralize pathogen. In the first term of Eq. (13), *S* is the sensitivity of the innate immune system. This term ensures that glucose consumption by the innate immune system is correlated with its activity. In this model, when innate immune cells desensitize due to anti-inflammatory signaling, they also release resources by stopping glucose uptake.

For the adaptive immune compartment, we model the glucose dynamics as follows:

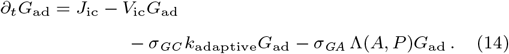

Thus, in addition to the baseline consumption rate *V*_ic_ as before, adaptive immune cells also use glucose to produce cytokines and consume more glucose when they proliferate. The parameter *σ*_*GA*_ controls how much glucose adaptive immune cells consume for the purpose of proliferation.

## Results

### Glucose dynamics upon immune response to infection and its role in pathogen clearance

We will begin by discussing a typical course of the immune response over time and the corresponding metabolic state of the individual in response to an infection [Fig. 4]. Shortly after infection, replicating pathogens such as bacteria or virions, or the amount of infected tissue, grow exponentially [Fig. 4A, logarithmic vertical axis]. In the absence of an immune response, pathogen would saturate at the carrying capacity.

**Fig. 4:**
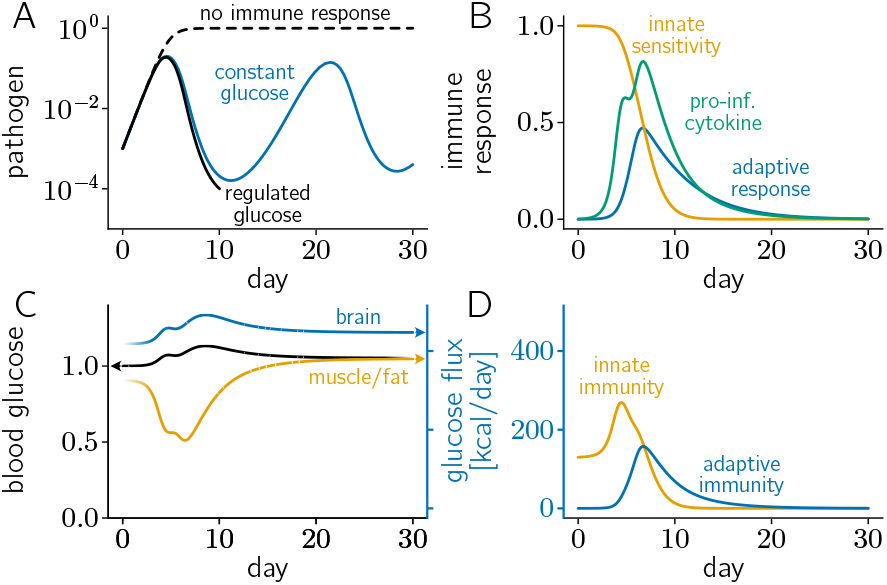
Typical within-host dynamics of the pathogen, the immune response, and the impact on glucose regulation. The species concentrations in Panels A-C are normalized as described in the text. A) The blue curve shows the pathogen dynamics if glucose levels were clamped at their homeostatic values. If glucose levels are regulated by the immune response, then pathogen is cleared (black curve). The immune response suppresses and eliminates replicating pathogen by B) secreting pro-inflammatory cytokines (green curve) and expanding adaptive immunity (blue curve). Cytokines are secreted by the innate immune system according to its sensitivity (yellow curve) to pathogen, which is initially high to detect pathogen and gradually diminishes to protect the host from excessive inflammation-induced damage, and by the adaptive immune system. C) Plasma and brain glucose levels increase during infection, whereas muscle and fat glucose levels decrease. Here, glucose levels are higher after infection was cleared than before, because the innate immune system has desensitized and reduced its glucose uptake. Over longer timescales, the sensitivity of the immune system would recover to *S* = 1 and the brain and plasma glucose levels would return to their homeostatic values. Note that this panel has a double axis. The right axis is synchronized with the axis of panel D, as indicated by its color code. D) Metabolic costs of activating the innate and adaptive immune responses. For simplicity, we neglected the production of antibodies; taking into account antibody production would make the immune response more robust. We also neglected the consumption of cytokines when clearing pathogen (*σ*_*CP*_ = 0), and the consumption of glucose when producing cytokines (*σ*_*GC*_ = 0), clearing pathogen (*σ*_*GP*_ = 0), and expanding adaptive immune cells (*σ*_*GA*_ = 0). All other parameter values are reported in the main text.

**Fig. 5:**
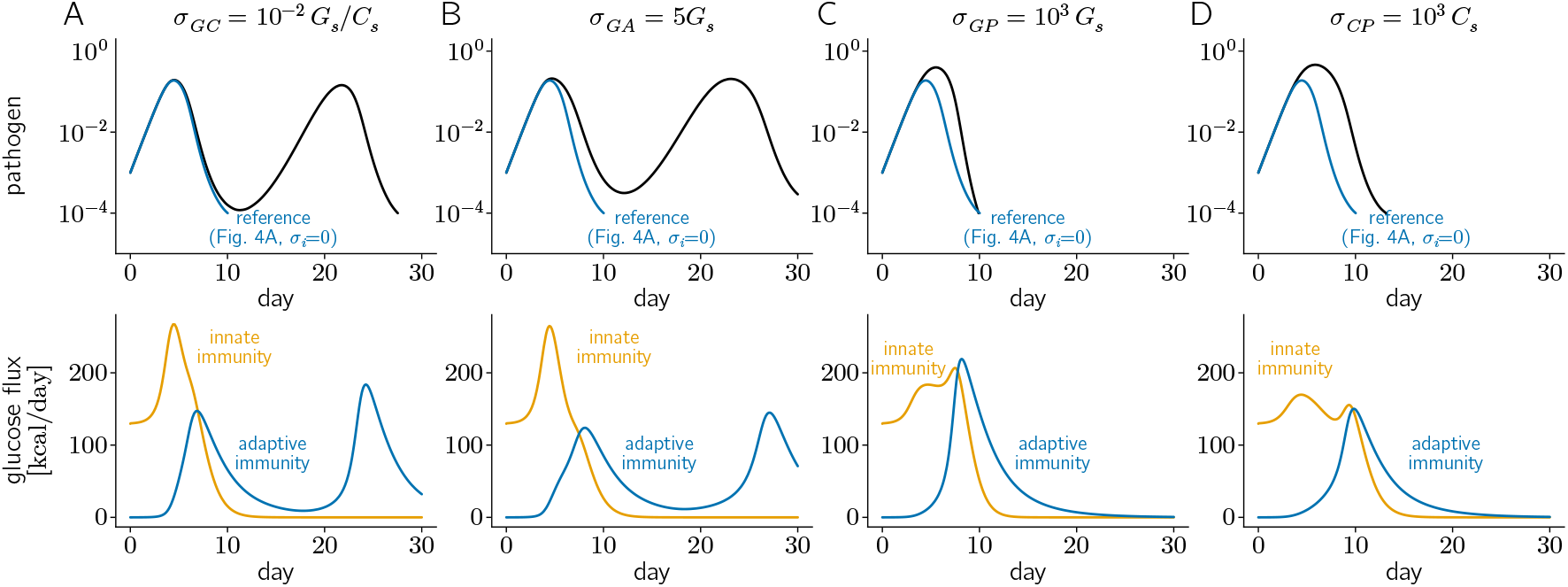
Comparison of reference simulations [black line in Fig. 4A] to scenarios where (A) glucose is explicitly consumed to generate cytokines, (B) glucose is consumed to expand adaptive immune cells, (C) glucose is consumed to eliminate pathogen, and (D) cytokines are consumed to eliminate pathogen. Top row depicts pathogen levels, with the blue curves representing the reference case (Fig. 4) and the black curves representing the dynamics with parameters changed as indicated. Bottom row depicts glucose flux into the innate and adaptive immune compartments. (A) Host outcomes are sensitive to increasing glucose consumption when producing cytokines, which can lead to a flare-up of infection before it is cleared, or more predator-prey cycles. Such predator-prey dynamics can also be seen in panel B, where we have strongly increased *σ*_*GA*_. (B-D) In contrast, the dynamics are much less sensitive to increasing glucose consumption when expanding adaptive immunity or killing pathogen, or cytokine consumption when eliminating pathogen. Better host outcomes correlated with larger glucose consumption by adaptive immunity. Parameter values are reported in the main text and in the panels.

**Fig. 6:**
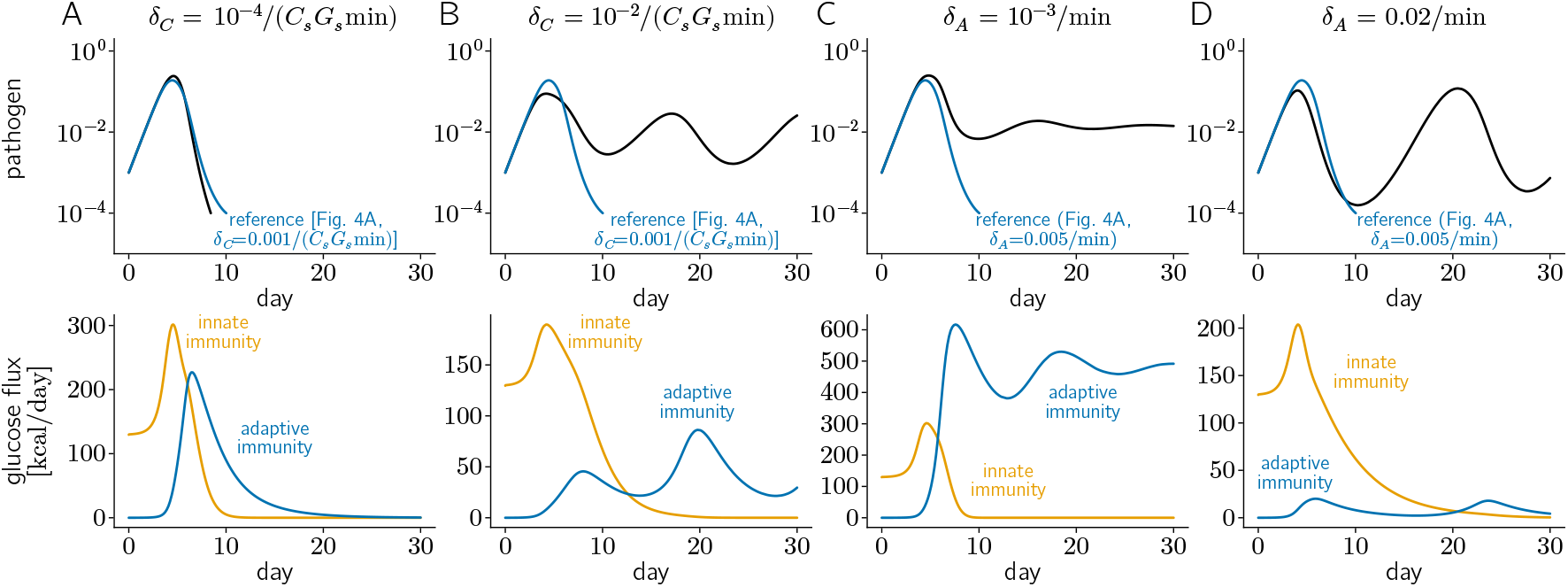
Comparison of reference simulations [black line in Fig. 4A] to scenarios where (A) innate immunity is less effective at killing pathogen, or (B) innate immunity is more effective at killing pathogen, or (C) adaptive immunity is less effective at killing pathogen, or (D) adaptive immunity is more effective at killing pathogen. Top row depicts pathogen levels, with the blue curves representing the reference (Fig. 4) and the black curves representing the dynamics with parameters changed as indicated. Bottom row depicts glucose flux into the innate and adaptive immune compartments. (A, B) Our simulations indicate that host outcomes are better, from the perspective of eliminating pathogen, when the innate immune response has a lower rate of pathogen clearance (panel A), because this is associated with a stronger adaptive immune response (blue curves in the lower row). (C, D) Comparing the pathogen dynamics (black curves) with the reference (blue curve) in the upper row suggests that there is an intermediate rate of pathogen clearance by adaptive immunity which optimizes host outcomes. The lower row shows that if the rate of pathogen clearance by adaptive immunity is too low, then it partially compensates with an increased metabolism but fails to eliminate pathogen. If the rate of pathogen clearance is high, then the energy demand of the adaptive immune response is reduced and pathogen rebounds. Parameters are reported in the main text and in the panels.

The innate immune response is the first line of defense of the host against the replicating pathogen. In response to the exponentially growing pathogen, cells of the innate immune system secrete pro-inflammatory cytokines until they begin to desensitize around days 3 to 4 [53], and inflammation is strong enough to suppress infection by inhibiting pathogen growth [Fig. 4A,B]. To metabolically feed the immune compartment, cytokine signaling inhibits pancreatic insulin secretion and insulin-dependent glucose uptake of muscle and fat tissues. The first effect leads to an increase in hepatic glucose production which, together with the reduced glucose uptake by muscle and fat tissues, increases plasma glucose levels [Fig. 4C].

Indeed, infections are known to cause hyperglycemia via the secretion of stress hormones and cytokine signaling [54, 55, 56]. The increased plasma glucose levels meet an increased demand from immune cells, which up-regulate the number of glucose transporters in response to cytokine signaling. Plasma glucose levels are highest when the metabolic cost of the innate immune response is maximal [Fig. 4D]. The associated metabolic rate is roughly consistent with estimates that the total energy demand of all leukocytes increases from 380 kcal*/*d at homeostasis to 500 kcal*/*d when they are activated [57]. Brain glucose levels also increase [Fig. 4C] as these levels track plasma levels [Fig. 3]. Although we do not observe hypoglycemia for the parameters chosen in Fig. 4, we note that an increased metabolic demand of the immune system, potentially due to severe sepsis [58], can have such an effect [cf. Fig. 7D].

**Fig. 7:**
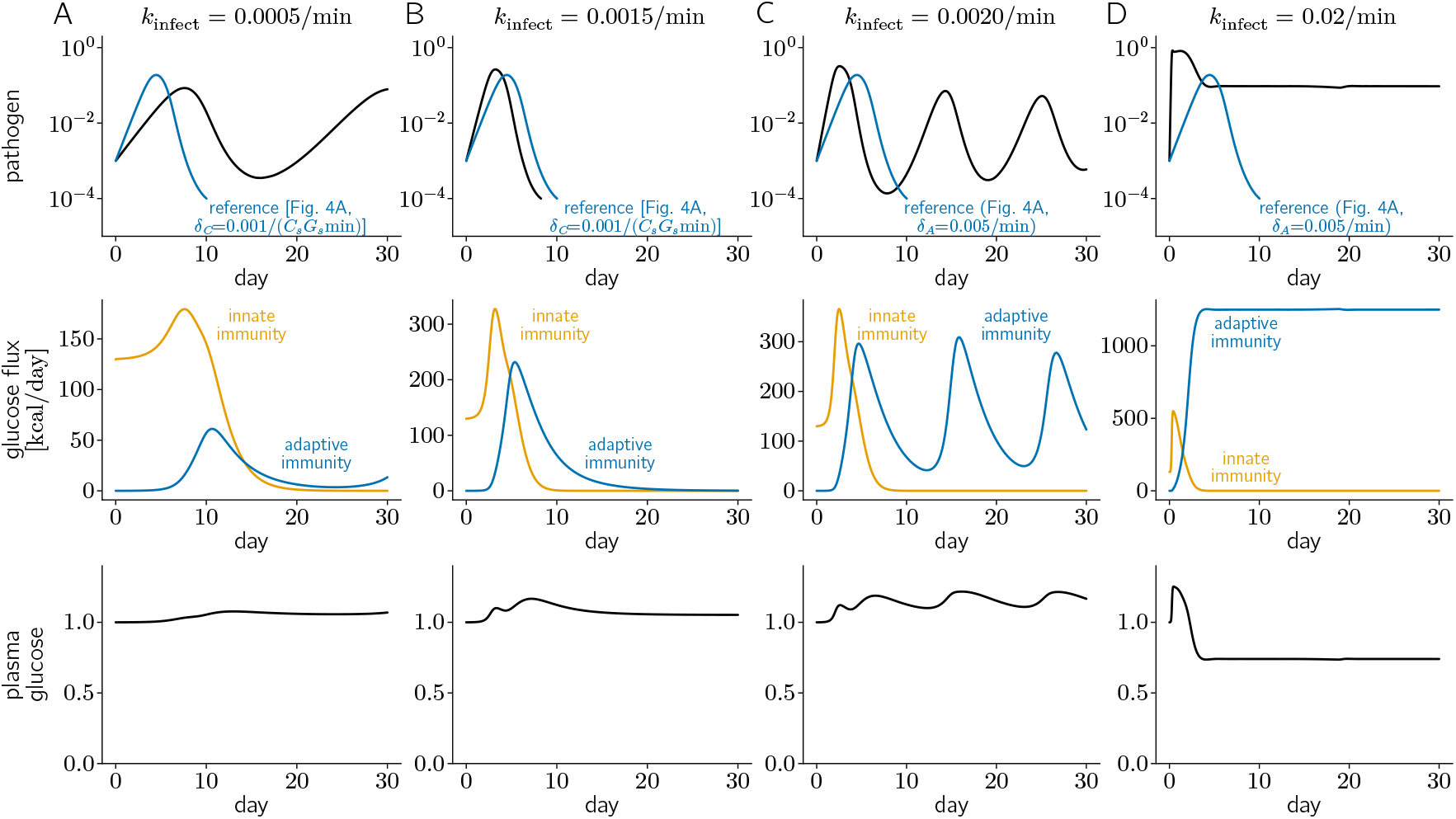
Comparison of reference simulations [black line in Fig. 4A] to scenarios where (A) pathogens replicate more slowly, or (B-D) pathogens replicate more quickly. Top row depicts pathogen levels, with the blue curves representing the reference (Fig. 4) and the black curves representing the dynamics with parameters changed as indicated. Middle row indicates glucose consumption by immune cells, and bottom row shows plasma glucose levels. (A) Slower pathogen replication can hide it from the immune response, thus causing a rebound of infection. Faster pathogen replication can either (B) improve host recovery, (C) lead to periodic flare-ups through predator-prey oscillations, or (D) cause persistent infection at constant pathogen levels. In the last scenario, for very high pathogen replication rates, the immune system consumes a large amount of energy which leads to a decrease in plasma glucose levels. Parameters are indicated in the main text and in the panels.

We tested how well the host can fight infection if the immune system does not regulate plasma glucose fluxes via cytokine signaling. To that end, we keep the plasma glucose levels fixed at *G* = 1 *G*_s_. Under these conditions, we find that the immune system fails to drive the pathogen to extinction. This is followed by a rebound of pathogen [blue line in Fig 4A], leading to another expansion of the adaptive immune response that drives pathogen levels down. These oscillations are reminiscent of predator-prey dynamics, akin to the immune response preying on pathogen. When pathogen levels decrease, so does the activation level of the immune response, which leads to an increase in the pathogen replication rate and exponential growth. This happens when the rate of pathogen clearance at peak immune activity is not high enough to cross the extinction threshold. This would be the situation if, for example, glucose fluxes were not regulated by cytokines, thus leading to inadequate energy resources for the immune system.

In addition to directly suppressing infection, cytokine-mediated inflammation also triggers the adaptive immune response by facilitating immune cell differentiation and growth and by increasing glucose uptake into immune cells. By the time the innate immune response has declined, the adaptive immune response is in full swing and takes over the host defense. In the ideal case, the adaptive immune response eliminates pathogen from the host. How does this outcome change (a) when innate or adaptive immunity have an increased or reduced ability to clear pathogen, (b) when the metabolic costs of producing cytokines, expanding the adaptive immune response, and clearing pathogen are high, (c) for pathogens with higher or lower replication rates, or (d) when immune control of host metabolism is perturbed? In the following, we answer these questions with numerical simulations of our model equations.

### Activity-dependent glucose and cytokine consumption

So far, we have neglected the task-specific consumption of cytokines and glucose when innate and adaptive immune responses eliminate pathogen. Moreover, we have also neglected the task-specific consumption of glucose when immune cells produce cytokines and when adaptive immune cells expand. In this section, we include these effects and study the resulting consequences for host outcomes. Our simulations show that increasing the number of units of glucose used to produce a single unit of pro-inflammatory cytokines, *σ*_*GC*_, has a detrimental effect on the host. Specifically, even fairly low values of *σ*_*GC*_ lead to a failure of the immune response to eliminate the pathogen. This is because, according to Eq. (13), increasing *σ*_*GC*_ will lead to glucose depletion in the innate immune compartment, thus limiting cytokine production. Similarly, according to Eq. (14), it will also drain glucose from adaptive immune cells, thereby inhibiting their replication. Both of these effects will impede immune function, suggesting that glucose levels should be sufficiently high to not be a bottleneck for cytokine production.

In contrast, the dynamics are much less sensitive to increasing glucose utilization for the expansion of adaptive immunity, *σ*_*GA*_. Only at high values of *σ*_*GA*_ is there a detrimental effect on the host. This is because somewhat higher levels of *σ*_*GA*_ effectively reduce the rate of expansion of adaptive immunity, but do not change the fact that it still grows exponentially. This is in contrast with the strong effect of *σ*_*GC*_ which deprives the innate immune response of glucose and leads to a failure to secrete cytokines, thereby inhibiting the ability to slow pathogen growth and activate adaptive immunity.

Similarly, increasing glucose and cytokine consumption during pathogen removal has no significant effect on host outcomes even at very high values of *σ*_*GP*_ and *σ*_*CP*_. In summary, the ability of the host to clear pathogens is predominantly sensitive to the amount of glucose that is consumed per unit of cytokines produced.

### Perturbations of the immune response

In this section, we study how changes in the rate constants corresponding to pathogen clearance by innate immunity and adaptive immunity affect host outcomes. Decreasing the rate of cytokine-dependent removal of pathogen by the innate immune system at early times [*δ*_*C*_ in Eq. (6)] counter-intuitively leads to faster infection clearance [Fig. 6A]. Increasing *δ*_*C*_ leads to persistent infection with predator-prey oscillations indicating a periodic flare-up of infection [Fig. 6B]. In fact, our model would predict that the best scenario for the host arises when innate immunity does not participate in pathogen clearance but solely acts to activate adaptive immunity, unless *δ*_*C*_ is large enough so that pathogen levels monotonically decay after inoculation at *t* = 0. This counter-intuitive observation can be explained as follows. Because inflammation caused by innate immune responses causes cell death in healthy host tissues, it must be self-limiting to prevent the innate immune response from excessively damaging the host. For example, negative feedback through the secretion of anti-inflammatory cytokines [Eq. (10)], tempers the innate immune response [45]. So, the innate immune response shuts off after a short duration. Thus, the removal of pathogen after it has a chance to grow can only be accomplished by an immune response that acts over time scales longer than the duration of the innate immune response; viz., the adaptive immune response. If the innate immune response is overly effective at clearing pathogen (unless it is sufficiently large—see above), it does not prevent pathogen growth, but limits it. This prevents the efficient activation of the adaptive immune response for reasons noted below.

By reducing pathogen load, high rates of cytokine-dependent removal of pathogen by the innate immune system also inhibit pathogen-stimulated cytokine release and thus lead to low cytokine levels. In turn, low cytokine levels lead to a low influx of adaptive immune cells through differentiation and chemotaxis. Low cytokine levels also lead to low glucose levels in immune cells, thus depleting a resource that is required for the expansion of adaptive immunity. Low pathogen load due to high rates of cytokine-dependent removal of pathogen by innate immunity also reduce the encounter rate of adaptive immune cells with antigen. This delays the expansion of adaptive immunity according to Eq. (8). These three effects lead to the inability of the adaptive immune response to decrease pathogen levels below the extinction threshold, resulting in rebound of the infection. Thus, an innate immune response that is overly effective at clearing pathogen, but not potent enough to eliminate pathogen by preventing its growth altogether (including at *t* = 0), would lead to inability to clear pathogen and cause greater immunopathology (which we have not explicitly modeled). This result may be related to the observation that a strong innate immune response or early initiation of antiviral treatments can result in inability to completely clear virus and cause later viral rebound in some people infected with SARS-CoV-2 [59]. Our results thus suggest a reason not noted before for why there is such tight control of inflammation through secretion of anti-inflammatory cytokines shortly after the activation of innate immune responses.

Our model neglects that small values of *δ*_*C*_ can be detrimental to the host for a different reason, namely morbidity induced by high pathogen and consequently high cytokine loads. In a realistic setting, the adaptive immune response takes about a week to produce effective antibodies, and a lacking innate immune response would mean uncontrolled pathogen growth. These findings suggest that there is a trade-off in choosing the potency of the innate immune response in clearing pathogen (parameterized by *δ*_*C*_).

Adaptive immunity is also ineffective when it eliminates pathogens too slowly, thereby maintaining persistent infection [Fig. 6C], or too quickly, which leads to periodic flares of infection. This is because low pathogen levels due to overactive adaptive immune responses inactivate these responses quickly, thus preventing extinction of the pathogen. The pathogen can subsequently rebound, and this cycle continues as in predator-prey dynamics [Fig. 6D]. However, we do not explicitly model infection of cells by pathogen or account for neutralizing antibodies that can prevent infection of new cells. The killing of pathogens embodied in Eq. (6) best represents elimination of infected cells by effector T cells. Neutralizing antibodies produced during the adaptive immune response can continue to protect against new infections even when effector T cells have stopped eliminating infected cells, thus driving the pathogen to extinction. This result shows the critical importance of antibodies in mediating effective adaptive immune responses.

### Effects of pathogen replication rate

In this section, we study how changes in the rate of pathogen replication affect host outcomes. Our first prediction is that slow pathogen replication can effectively hide the infection from the adaptive immune response, leading to a rebound at later times [Fig. 7A]. This observation has the same underlying reason as the worse host outcomes when the innate or adaptive immune responses are too effective at clearing pathogen. However, high pathogen replication rates are also detrimental to the host and can lead to persistent infection or host death [Fig. 7C,D]. Depending on the parameters, persistent infection is characterized by pathogen loads that grow exponentially until they reach carrying capacity (not shown), by oscillations with periodic flare-ups due to the predator-prey dynamics discussed previously [Fig. 7C], or by relatively low pathogen loads that are kept in check by a highly active immune response [Fig. 7D]. The last scenario arises for very large pathogen replication rates, which lead to high cytokine levels and activity of the adaptive immune response. Together, these immune responses keep pathogen levels low despite a high replication rate, but at a high metabolic cost. Interestingly, in this parameter regime the plasma glucose levels drop below homeostatic concentrations. Clinically, a drop in plasma glucose levels towards hypoglycemia is very rare in sepsis but associated with an especially grim prognosis [58], which is consistent with fast pathogen replication and cytokine storm.

## Discussion

In this paper, we have studied the dynamical interplay between the immune responses to infection and regulation of redistribution of glucose resources to different tissues. Toward this end, we developed a systems-level physiological model. Our results have implications for how mis-regulation of the interplay between the role of immunity in clearing infection and mediating adaptation of the redistribution of metabolic resources could result in detrimental effects for the host. For example, we find that cytokine-mediated redistribution of glucose resources is very important for effective clearance of infection. Clamping the glucose level to the homeostatic value results in ineffective pathogen clearance. Also, too high a consumption rate of glucose by innate immune cells to carry out effector functions inhibits the ability to effectively clear infection. This is because a high rate of glucose consumption limits cytokine production too soon, thus inhibiting various immune functions. We also study how the strengths of the innate and adaptive immune systems and pathogen replication rates affect metabolic adaptation and clearance of infection. Three examples of such findings are: (1) Unless the innate immune response is strong enough to eliminate the pathogen, too strong an innate immune response can lead to very low pathogen levels at early times, which inhibits activation of the adaptive immune response and effective clearance of infection. In addition to limiting immunopathogenesis, this may be an additional reason for limiting the inflammatory response through the secretion of anti-inflammatory cytokines. (2) Very low pathogen replication rates allow the pathogen to “hide” from the immune system and rebound at later times. (3) While pathogens with very high replication rates can be controlled to low levels, the metabolic cost to the host becomes very large. High immune activity due to a large pathogen replication rate, such as during sepsis and cytokine storm, could lead to hypoglycemia.

**Table 1.**
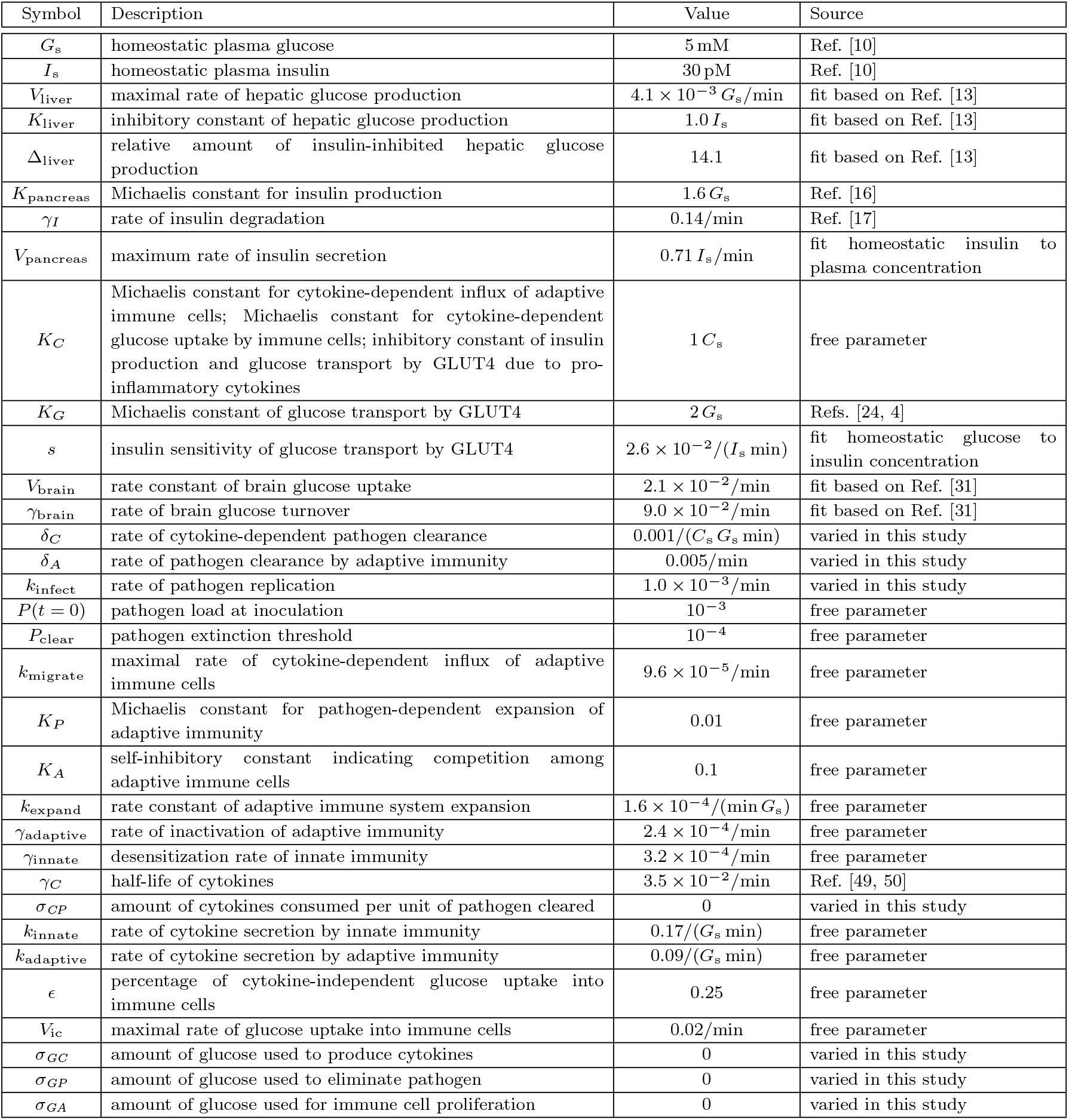
Table of parameters used in this study, indicating which ones are known or fitted from prior literature, and which ones are unknown with reasoning provided in the main text.

In our model, the physiological system seeks to restore a homeostatic state via an array of negative feedback loops, such as: enhanced plasma glucose leads to insulin secretion which reduces glucose, pathogen replication leads to an immune response which clears pathogen, and inflammation during the immune response leads to secretion of anti-inflammatory molecules which reduce inflammation. The crosstalk between these different elements is also based on negative regulation, whereby cytokines inhibit insulin production and insulin-dependent glucose uptake, thus effectively increasing plasma glucose levels. We speculate that the many mechanisms for negative regulation could be beneficial for system stability and robustness in returning the system to homeostasis after infection. In contrast, the immune response itself features positive feedback loops, such as between the expansion of adaptive immune cells and cytokine secretion. Such positive feedback, together with exponential growth of the adaptive immune compartment, could be useful to outgrow and ultimately eliminate replicating pathogen.

One consequence of negative regulation is an undershoot before reaching a steady state. This can be experienced, for example, during hypoglycemia after consuming large amounts of carbohydrates. This feature is also present in the interactions between pathogens and the immune system discussed here and falls into the category of predator-prey systems known from population dynamics, which can show oscillations. The corresponding undershoot is useful for clearing pathogen by crossing the extinction threshold. At the same time, it also leads to counter-intuitive effects, which are rooted in the fact that predator-prey oscillations are abolished when there is a separation of timescales between prey (pathogen) and predator (immune system) dynamics. Thus, reducing the undershoot by clearing pathogen too quickly or too slowly can lead to persistent infection. In other words, there is an optimal pathogen clearance rate that improves host outcomes. Conversely, pathogens can potentially escape the immune response by up-or downregulating their replication rate. The latter scenario could apply, for example, to tuberculosis or leishmaniasis.

To combat infection, the body diverts resources to the immune system in the form of glucose. This is achieved by increasing plasma glucose concentration via two mechanisms: reducing insulin which in turn leads to larger hepatic glucose production, and in addition directly inhibiting insulin-dependent glucose uptake into muscle and fat tissues. Having increased the glucose supply, pro-inflammatory cytokines also lead to incorporation of GLUT1 into the plasma membrane of immune cells, thereby upregulating their glucose uptake. For typical immune activity, these effects together increase plasma glucose levels and brain glucose consumption. This is consistent with the simulations of Zhao, Straub, and Meyer-Hermann [5], but arises in their model due to a different mechanism. In their work, this can be mainly attributed to an increase in lipolysis by immune activity, which is conceptually analogous to the increase in hepatic glucose production due to the inhibition of insulin by cytokines, which we described. Brain glucose levels increased in Ref. [5] due to a larger plasma glucose concentration despite a decrease in the brain glucose uptake rate due to immune activity. Similarly, metabolic fluxes towards the immune system also increased in Ref. [5] due to larger plasma glucose values despite an auto-inhibition of the glucose uptake rate by the immune system. This is akin to the anti-inflammatory effect mediated by the secretion of anti-inflammatory cytokines which we have taken into account. Thus, there can be multiple pathways that ultimately lead to similar phenomenology.

In this paper, we have studied the interplay between adaptation and immunity in the context of adaptation of metabolic (glucose) fluxes upon infection. However, some of our conclusions may be generally applicable to many other situations where the interplay of the dynamics of adaptation of a limiting resource and the immune response plays a role in maintaining health or mediating disease after physiological perturbations.

